# A machine learning approach for MRI-based classification of individuals with mild cognitive impairment

**DOI:** 10.1101/2021.05.27.445930

**Authors:** Roni Tibon

## Abstract

Diagnosis criteria for Mild Cognitive Impairment (MCI) rely on cognitive symptoms and do not require specific biomarker evidence of pathology. Nevertheless, patients may still have subtle brain changes that are identifiable with neuroimaging. To classify MCI patients vs. healthy age-matched controls, two algorithms—Multilayer Perceptron (MLP) and Support Vector Machine (SVM)—were applied to structural MRI data from the OASIS-1 cohort (https://www.oasis-brains.org/). Despite their comparable performance in some measures, the MLP algorithm was preferable due to superior recall of MCI cases.

## 1 Introduction

Alzheimer’s disease (AD) is an age-related neurodegenerative disorder, affecting millions of people world-wide. It is characterised by progressive dementia, from mild memory impairment to global cognitive dysfunction and eventually death^1^. Although there is currently no cure for AD, early detection enables effective management of the disease, as well as the ability to prevent or delay dementia.

Mild Cognitive Impairment (MCI) is a syndrome defined as cognitive decline greater than expected for an individual’s age and education level but that does not interfere notably with activities of daily life^2^. Although the diagnosis criteria for MCI rely solely on cognitive symptoms, and do not require specific biomarker evidence of underlying AD pathology^3^, patients may still have subtle brain changes that are identifiable with neuroimaging. Importantly, MCI is commonly a prodromal state of AD, with a high probability of progression to dementia^4^. Therefore, identifying reliable biomarkers for detection of MCI, can be a step towards early detection and prevention of AD.

### 1.1 The OASIS Dataset

The Open Access Series of Imaging Studies (OASIS)^5^ is a series of publically available datasets of magnetic resonance imaging (MRI) of the human brain. For each individual in these datasets, T1-weighted MRI scans were obtained. The release used for the current project includes cross-sectional data from 416 right-handed, male and female individuals across the adult life span age 18 to 96 years. The dataset (as posted on Kaggle: https://www.kaggle.com/jboysen/mri-and-alzheimers) includes 3 neural measures:

- Atlas Scaling Factor (ASF): Computed scaling factor that transforms native-space brain and skull to the atlas target (i.e., the determinant of the transform matrix). This is a continuous measure that ranged between 0.88 and 1.56 (M=1.22, SD=0.13) in the current sample.
- Estimated total intracranial volume (eTIV): Automated estimate of total intracranial volume in native space derived from the ASF. This is a continuous measure that ranged between 1123 and 1992 (M=1461.4, SD=162.1) in the current sample.
- Normalised whole-brain volume (nWBV): Automated tissue segmentation based estimate of brain volume (gray-plus white-matter). Normalised to percentage based on the atlas target mask. This measure is continuous and ranges between 0 and 1. Range in the current sample ranged between .64 and .85 (M=.75, SD=.05).

The dataset further includes each individual’s score in two measures of clinical assessment. The one used here is the Mini–Mental State Examination (MMSE): a 30-point questionnaire commonly used to measure cognitive impairment^6^. The scale is discrete (ranges between 0 and 30), with any score of 24 or more indicating normal cognition, scores between 20 and 23 indicating mild cognitive impairment (the diagnostic criteria for MCI), and scores below 20 indicating moderate or severe cognitive impairment. The range in the current sample (after preprocessing, see below) was between 20 and 30 (Med=29, IQR=27-30). MMSE scores were reported for a subset of 235 individuals; only this subset was included in the current study. Additional demographic details (gender, socio-economic status, and education level) were also available, but were not used here.

### 1.2 Algorithms

#### 1.2.1 Multilayer Perceptron (MLP)

The multilayer perceptron (MLP) is a supervised learning algorithm. It consists of a system of interconnected nodes, which can represent a nonlinear mapping between an input vector and an output vector. In this feed-forward system, inputs for each node are multiplied by the weights in a node, summed together, transformed via an activation function which defines the specific output or “activation” of the node, and fed forward to be an input to the nodes in the next layer of the network. The backpropagation algorithm is commonly used for training. In short, to train an MLP via backpropagation, the network weights are initialised and a training input vector is propagated through the network to obtain an output. The error signal is then calculated by comparing the obtained output with the target output, and is propagated back through the network, and the weights are repeatedly adjusted to minimise the error^7,8^. It has been shown^9^ that the multilayer perceptron can be trained to approximate any smooth, measurable function, including highly non-linear functions. Unlike other statistical techniques, MLP makes no prior assumptions concerning the data distribution, and can be trained to accurately generalise when presented with new, unseen data. One limitation of MLP (and neural networks in general), is that it is difficult to interpret. In addition, the performance of the backpropagation algorithm is greatly affected by data dimensionality. That is, as the dimensionality of the data increases, the amount of training data required by the MLP rapidly increases. Similarly, as the data become more complex the number of weights, and hence the size of the network, rapidly increase. Both these problems serve to slow the speed at which the algorithm will converge.

#### 1.2.2 Support Vector Machine (SVM)

A support vector machine (SVM) is a supervised machine learning model that finds the decision boundary for which the functional margin between two data classes is maximised. In order to apply SVM to problems that are not linearly separable, the data from the original space is transformed into a higher dimensional feature space. The “kernel trick” is applied to support the transformations that include many data features and many polynomial combinations of these features in a relatively low computational cost. SVM is considered an efficient and versatile algorithm, which is effective in high-dimensional space. Its main drawback is that unlike MLP, it does not directly provide probability estimates (though those can be calculated via cross-validation). Moreover, additional steps should be taken when the number of features is much greater than the number of cases in order to avoid overfitting^[7]^.

### 1.3 Hypothesis Statement

MLP usually outperforms SVM against the same dataset, when the number of features is much greater than the number of samples. Nevertheless, in the current study, the number of features was much lower than the number of samples. Moreover, the architecture of the MLP network was relatively shallow (with only one hidden layer), making it more comparable to SVM. Therefore, accuracy was expected to be similar across the two algorithms. In addition, MLP is generally more prone to overfitting and local minima trapping compared to SVM. Although several measures were applied to minimise these problems (e.g., defining momentum and restricting the number of epochs; see full details below), it is still possible that SVM will generalise to new data better than MLP. In terms of time complexity, because SVM models set their decision boundary on the basis of the sole support vectors, they are generally very fast to train, and so the training is expected to be quicker for SVM vs. MLP. However, given the limited number of data-points, the relatively simple architecture, and the low number of epochs, training time for MLP is also expected to be low.

## 2 Methodology

### 2.1 Data Preprocessing

Cases without MMSE scores were excluded. Remaining cases were classified according to their MMSE score: 0-mild cognitive impairment (MCI group; 19 < MMSE score < 24); 1-no cognitive impairment (Ctrl group; MMSE score >= 24); Additional cases, classified as severe cognitive impairment (MMSE score =< 19, N=12) were excluded, due to the current focus on early detection. Furthermore, in order to avoid confounding effects of age (which is also associated with structural changes of the brain), Ctrl group data were trimmed to include the same age-range as that included in the MCI group. Following these steps, the final dataset included 223 cases, 193 in the Ctrl group and 30 in the MCI group. All features were standardised by removing the mean and scaling to unit variance. Histograms of features distribution, boxplots of group data, and a correlation matrix of all variables included in the model, are shown in Figure 1.

**Figure 1.**
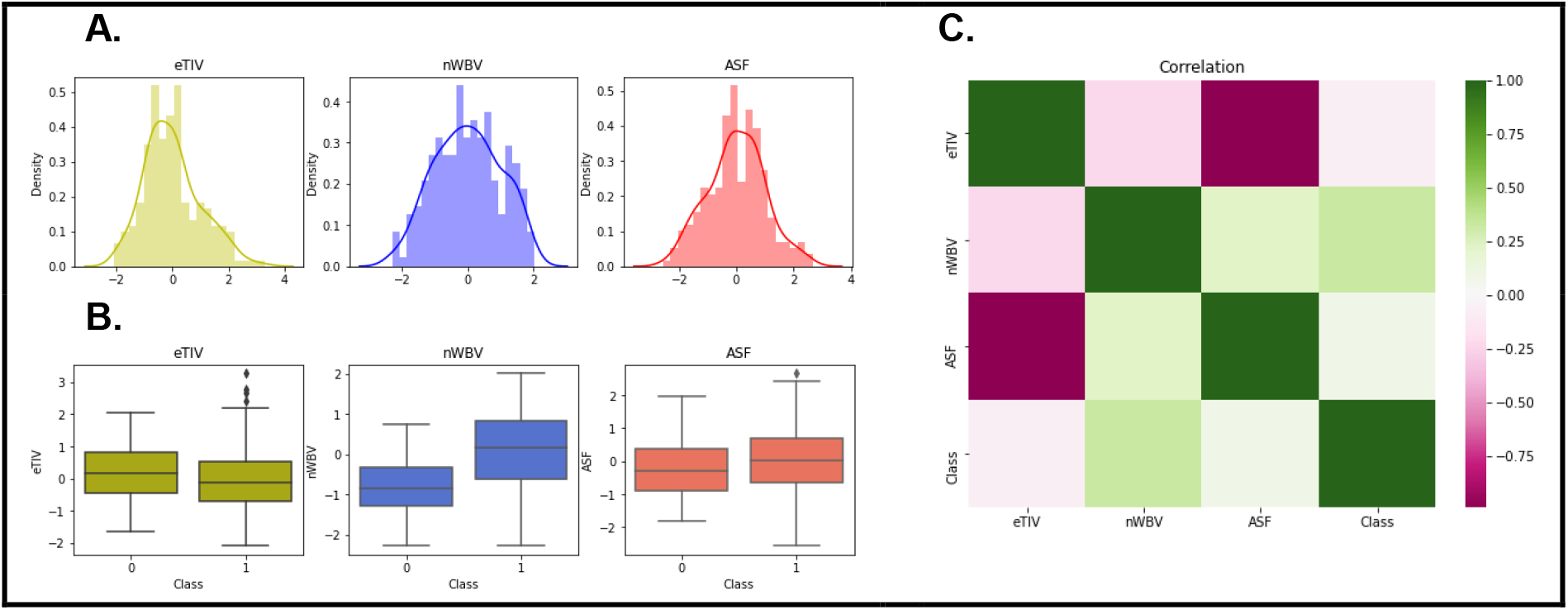
Properties of the data used in the study. (A) Histograms and density plots depicting the distribution of the preprocessed data, separately for each feature. (B) Boxplots depicting group data for the MCI group (Class 0) and the Ctrl group (Class 1) for each feature. (C) Correlation matrix depicting Pearson correlations between all variables included in the study.

### 2.2 Training and Evaluation

10% of the original (shuffled) dataset was held-out in order to be used as a test set for selected models. Because the dataset used in the current study is imbalanced, Synthetic Minority Oversampling Technique (SMOTE)^10^ was applied on the remaining 90%, in order to improve classification performance on the minority group. The transformed dataset was used for training and validation in the process of model selection.

An exhaustive grid search was employed to adjust the (hyper)parameters, separately for each algorithm. To select the best model for each algorithm (model selection), the models were evaluated using a stratified 10-fold cross-validation, in which the folds are selected so that the mean response value is approximately equal in all the folds. Model scoring was determined based on its accuracy. To compare the two algorithms (algorithm comparison), the selected (“best”) trained model for each algorithm was then evaluated against the entire train set and, more importantly, against the held-out test set.

### 2.3 Architecture and Parameters

Models were kept relatively simple due to the limited number of data points available in the current study. ***MLP models*** included three neurons in their input layer (3 features), two neuron in their output layer (for binary output), and one linear hidden layer comprised of a varying number of neurons (either 3 or 5). To allow non-linearity, non-linear activation was applied to the hidden layer after the linear transformation, using either ReLU or Leaky ReLU. Values obtained from the output layer were passed through the Softmax function in order to represent the model’s predicted probabilities for each class. Hyperparameters were selected to augment and control the optimization process. The batch size (either 16, 32, or 64) and learning rate (either 0.01, 0.1 or 0.5) were tuned to increase model stability and learning speed. The maximum number of epochs was set to 10, and early stopping was allowed, as preliminary investigation (see Implementation Details) suggested that the models start overfitting early on. A Stochastic Gradient Descent (SGD) optimization algorithm (with momentum set to 0.7, to avoid local minima) was used to backpropagate the prediction loss and adjust the parameters by the gradients. Thus overall, the MLP grid search included 36 parameter combinations: 2 [number of neurons in the hidden layer] X 2 [non-linear activation functions] X 3 [batch sizes] X 3 [learning rates].

For ***SVM models***, C-Support Vector Classification (SVC) was used. This algorithm maximises the margin while incurring a penalty when a sample is misclassified, or occurs within the margin boundary. The term C (which was set to either 0.01, 1, or 100), controls the strength of this penalty, thus acting as an inverse regularization parameter. Several kernel functions (linear, polynomial, radial basis function [RBF], and sigmoid) were tested, in order to allow non-linearity by mapping the training vectors into a higher dimensional space. Finally, gamma was set to either 0.0001, 0.01, or 0.1, to define how much influence a single training example has, with larger gamma indicating that other samples need to be closer in order to be affected. Overall, the SVM grid search included 36 parameter combinations: 3 [C] X 4 [kernels] X 3 [gamma] (though note that the actual space was in fact smaller, because gamma cannot be applied for a linear kernel; more details in the “implementation details” section below).

## 3 Analyses and Critical Evaluation

### 3.1 Model Selection

Figure 2 depicts the cross-validation accuracy (ordered from highest to lowest) for the MLP and the SVM models, following a grid search with various combinations of (hyper)parameters (see above). The standard deviation of the validation accuracy across the various models was greater for SVM than for MLP (STD_SVm_M_ = 0.13 vs. STD_MLP_M_ = 0.06). In contrast, within each model, the mean standard deviation across folds (n=10) was slightly greater for MLP than for SVM (STD_SVM_F_ = 0.07 vs. STD_MLP_F_ = 0.08). Taken together, this suggests that SVM is more influenced by specific parameter combinations and should therefore be tuned with cautious, whereas MLP is more sensitive to the particular selection of training samples, which might affect the model’s ability to generalise.

**Figure 2.**
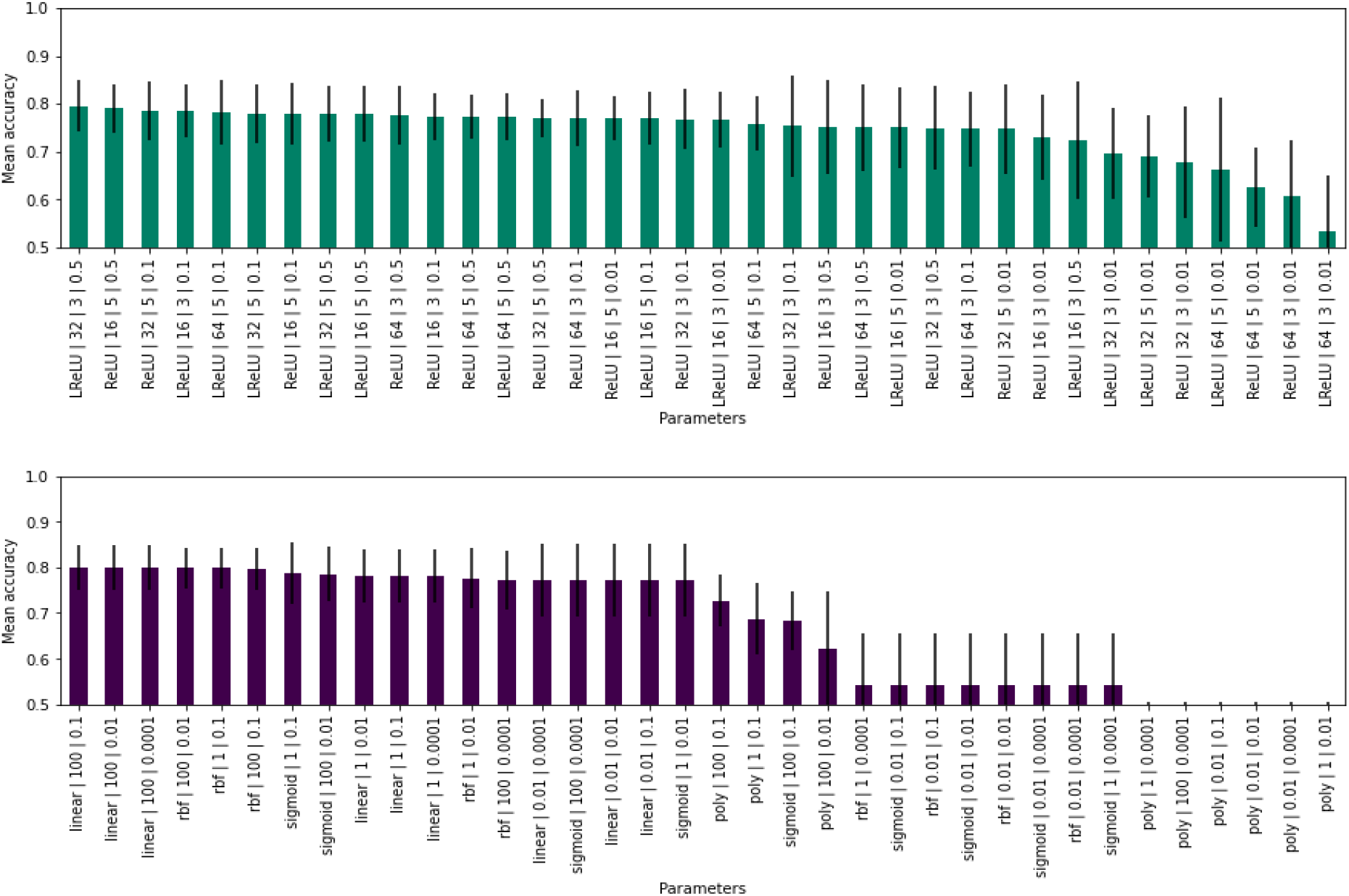
Results of the model selection process. Cross-validation accuracy (ordered from highest to lowest) following a grid search with various combinations of (hyper)parameters is shown for MLP (top) and the SVM (bottom) models. For MLP, legends on the x-axis indicate [activation function | batch size | hidden units | learning rate]. For SVM, legends on the x-axis indicate [kernel | C | gamma].

The figure further suggests that for both MLP and SVM, models can be grouped into those with classification accuracy > 75%, for which accuracy is highly similar, and those with lower classification accuracy, for which accuracy is more diverse. For SVM, the latter group included models with polynomial kernel and low gamma, suggesting that these transformations do not aid the separation of the data. For MLP, the latter group mainly included models with low learning rate, possibly suggesting that the restricted number of epochs used here (to avoid overfitting) was not sufficient for adequate training at this slow pace.

### 3.2 Algorithm Comparison

For validation of the models against the training set, the best model for each algorithm was fit on the entire training set (which included 90% of the data). For validation against the test set, the best model was fit on the remaining held-out 10%. Confusion matrices for the validation against the training and testing sets, as well as the respective ROC curves, are shown in Figure 3, panels A and B. ROC curves were comparable across algorithms, with similar AUC scores for MLP and SVM for the train set (83% and 84%, respectively), and slightly higher scores MLP than SVM for the test set (62% vs. 59%, respectively). In terms of overall accuracy, SVM outperformed MLP for both the train set (80% vs. 79%) and the test set (61% vs. 57%).

**Figure 3.**
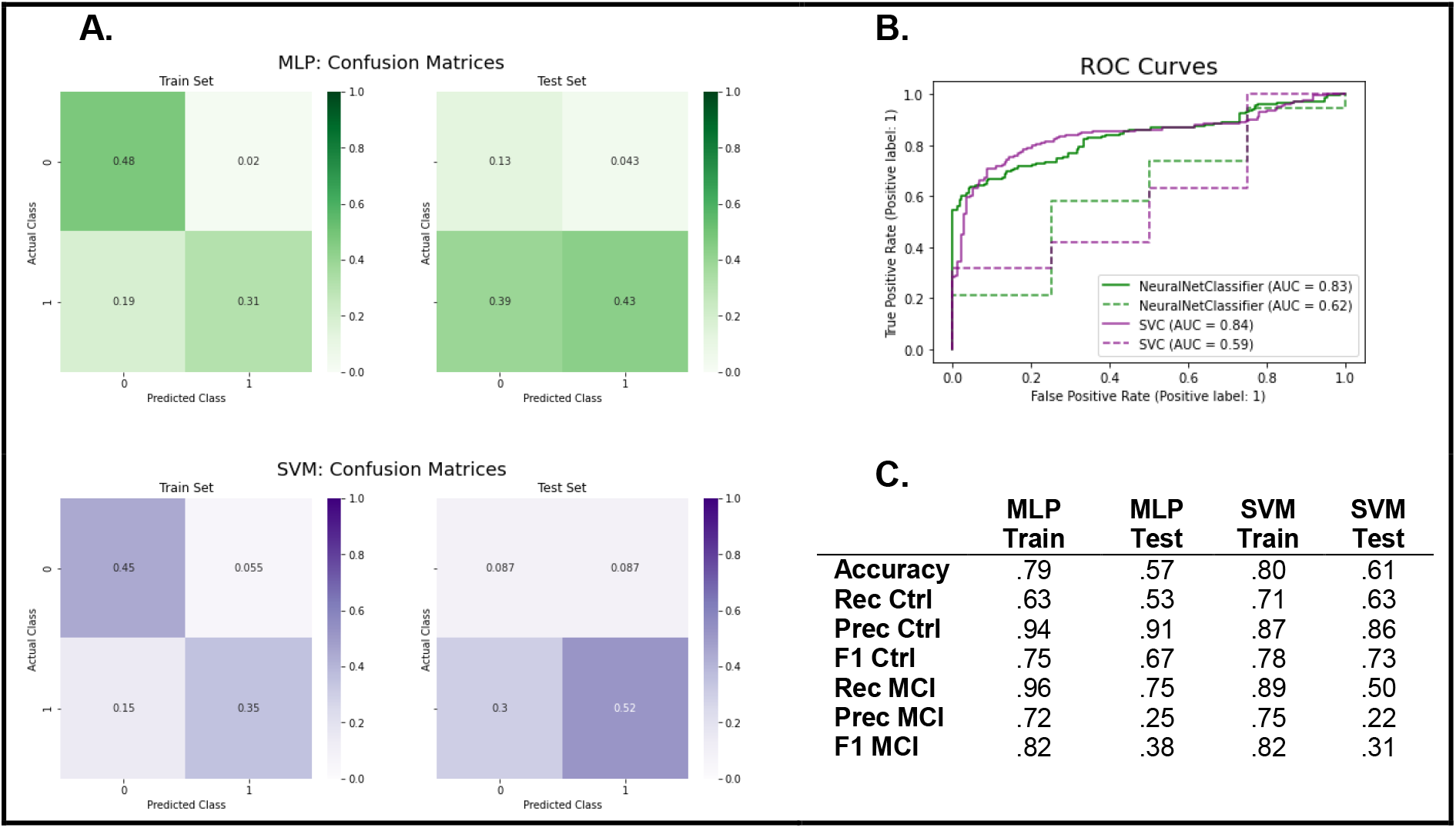
Results of the algorithm comparison process. Classes are 0-MCI, and 1-Ctrl. (A) Normalised confusion matrices for MLP (top) and SVM (bottom). On the left is the validation against the entire train set, and on the right is the validation against the (unseen) test set. (B) ROC curves for MLP (green) and SVM (purple), with the classification algorithm applied to the training set (solid lines) and test set (dashed lines). AUC scores are shown on the bottom-right. (C) Metrics used for the evaluation of the algorithms, including overall accuracy, as well as precision and recall reported separately for each group.

Importantly, however, as indicated by the confusion matrices, unlike the training set for which accuracy rates were similar for both classes, for the test set accuracy rates decreased significantly for the MCI group. This is likely the result of the imbalanced number of observations, with MCIs being a minority group. As SMOTE was applied to the train set, the distribution of cases was balanced for this set, resulting in similar classification accuracy for both groups. Nevertheless, the test set remained imbalanced, and therefore classification accuracy of cases belonging to the minority group was hinder. Importantly, even though classification accuracy for the minority group was low, it is still much better than what it was before SMOTE was applied (in which case, all cases were classified as controls, and none were classified as MCI; see Implementation Details). This suggests that applying some method for dealing with imbalanced data is necessary with the current dataset, and other methods (aside from SMOTE) should be explored in future studies.

Overall accuracy rates, as well as ROC curves and their associated AUC scores, can be misleading when applied in imbalanced classification scenarios^11^. Therefore, additional metrics—recall, precision, and F1 scores for the control and MCI group—were calculated and are shown in Figure 3, Panel C. For the train set, recall, precision, and F1 scores ranged between 63-96%. For Ctrl cases, recall and F1 scores were higher with SVM during both training and testing, whereas precision scores were higher for MLP. For MCI cases, recall scores were higher for MLP during both training and testing, precision scores were higher for SVM during training but for MLP during testing, and F1 scores were comparable during training but higher for MLP during testing.

Notably, the difference in recall scores for the MCI group is critical for the evaluation of the models. Namely, the main purpose of this study is to identify those who exhibit cognitive impairments, to aid early interventions and further evaluation. Therefore, the importance of correctly classifying Ctrl cases is inferior to the ability to accurately classify MCI cases. Recall of MCI cases, calculated as the fraction of MCI cases classified as MCI among all relevant elements, therefore represents the most important metric for model evaluation, and produces significantly better results for MLP than SVM (96% vs. 89% during training, and 75% vs. 50% during testing).

## 4 Conclusions

The aim of the study was to explore whether structural brain measures can be used to identify individuals exhibiting mild cognitive impairment, as indicated by their performance the mini mental-state examination (MMSE). Despite their comparable performance in some measures (e.g., in terms of accuracy), the MLP algorithm is concluded to be preferable. This is due to its superior recall of MCI cases—the main requirement from an algorithm aiming to aid early detection.

Nevertheless, in all measures, a sharp decrease in performance was observed when the models were applied to the held-out test set, suggesting low ability to generalise to new cases. While further tuning of the parameters might improve the performance of the models and should be explored, the models would probably benefit more from the inclusion of additional cases (especially in the MCI group) and additional features representing other structural properties of the brain. In particular, measures of specific brain regions, known to be affected by AD (e.g., the Hippocampus) should be included on top of the coarse, whole-brain measures that were used in the current study.

Before concluding, it is important to address the translational value of the study to real-life problems. An important question is whether or not the current approach is not only possible, but also viable. Or in other words – why bothering with the acquisition of costly brain images and the application of time-consuming ML algorithms, just to get the same results that a simple 10-item questionnaire can produce? The answer to that relies on the divergent between MMSE vs. ML based classifications: if, for example, cases classified as MCIs by both MMSE and ML would end up developing AD later on, while cases classified as MCIs by MMSE ***but not*** by ML would not, than the ML models provide valuable information, essential to the prognosis and care plan. While additional longitudinal data is required to allow such conclusions, the current study represents an important first step in this direction.

The code used for this project is available on: https://osf.io/msx4r/

## 6 Glossary

### Alzheimer Disease (AD)

Age-related neurodegenerative disorder, characterised by progressive dementia, at a level that interferes with daily tasks.

### Backpropagation

Widely used algorithm for training feedforward neural networks by computing the gradient of the loss function with respect to the weights of the network.

### Grid Search

Tuning technique that attempts to compute the optimum values of hyperparameters by performing an exhaustive search on the specific parameter values of a model.

### Kernel Trick

Algorithm that allows the transformation of data from the original space into a higher dimensional feature space by operating on an implicit space, without computing the actual coordinates of the data.

### Mild Cognitive Impairment (MCI)

Syndrome defined as cognitive decline greater than expected for an individual’s age and education level, but that does not interfere notably with activities of daily life. In many cases, MCI is a prodromal state of AD.

### Mini-Mental State Examination (MMSE)

24-item questionnaire commonly used to indicate the presence of cognitive impairment.

### Multilayer Perceptron (MLP)

Class of feedforward artificial neural networks used for supervised learning, consists of at least three layers of nodes: an input layer, a hidden layer and an output layer.

### Open Access Series of Imaging Studies (OASIS)

Series of publically available datasets containing MRI images of the human brain.

### Overfitting

Modelling error that occurs when a function is too closely fit to a limited set of data points, thereby reducing the model’s ability to generalise.

### Recall and Precision

Classification matrices used to assess the proportion of actual positives that were identified correctly (recall) and the proportion of positive identifications that were actually correct (precision).

### Receiver Operating Characteristic (ROC)

Curve – Graph showing the performance of a classification model at all classification thresholds. The **<u>Area Under the Curve (AUC)</u>** measures the two-dimensional area underneath the entire ROC curve.

### Rectified Linear Unit (ReLU)

Type of activation function, defined mathematically as y = max(0, x).

### Leaky ReLU

is similar, but has a small slope for negative values.

### Softmax

Function that converts a vector of numbers into a vector of probabilities ranging between 0 and 1.

### Support Vector Machine (SVM)

Supervised machine learning model that finds the decision boundary for which the functional margin between two data classes is maximised.

### Synthetic Minority Oversampling Technique (SMOTE)

Method for over-sampling the minority class, which involves creating synthetic minority class examples.

### T1-weighted MRI

Commonly use MRI sequence, used for the acquisition of structural brain images.

## 7 Implementation Details

### 7.1 Unbalanced Data

The figure below shows a reproduction of Figure 3, Panel A (confusion matrices) when SMOTE is not applied. As shown in the figure, for both SVM and MLP, without SMOTE the models predicted the majority class for all cases, thus render these models unusable.

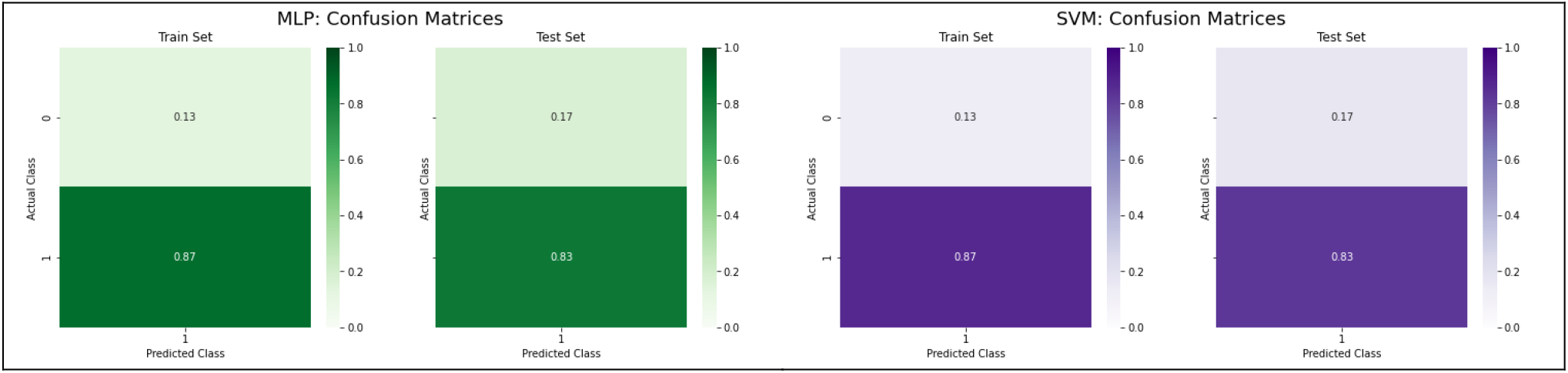

### 7.2 Number of Epochs

The number of epochs was initially set to 50 (without early stopping), and a figure displaying the loss for train and validation was computed for the best estimator (see below). On the left are the results of the preliminary implementation, with 50 epochs and no early stopping. On the right are the results of the final implementation with 10 epochs and early stopping. The figure suggested that the model converge and start overfitting early on, and so the maximal number of epoch was set to 10, and early stopping was applied.

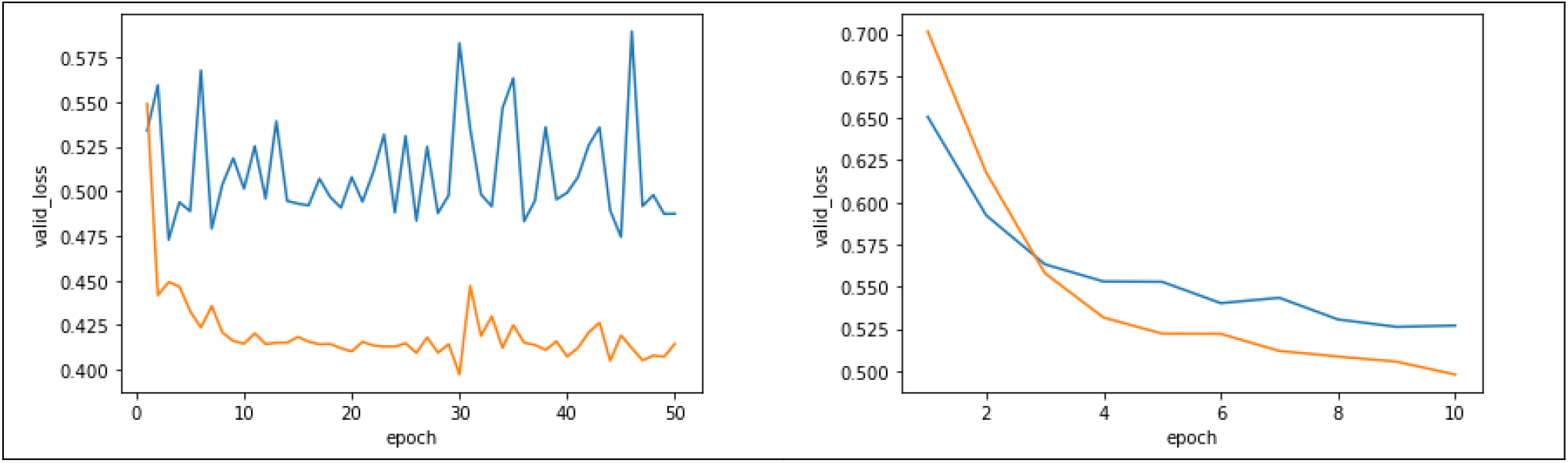

### 7.3 Additional Parameters

Aside from the parameters included in the grid search, which were tested systematically, some additional (hyper)parameters were tested manually during a preliminary investigation. These included (1) various dropout rates – all resulted in decreased performance relative to the fully connected model, and so dropout was not implemented in the final version; (2) replacing SGD with an Adam optimiser – resulted in reduced generalisation to the test set and was therefore excluded; and (3) additional variations (in addition to the ones reported in the paper) to the number of units in the hidden layer (ranging between 4 and 32) and inclusion of a second hidden layer (with varying number of units) – these also resulted in reduced generalisation to the test set. See comment in the notebook regarding these implementations.

## References

[1] Masters, C. L., R. Bateman, K. Blennow, and C. C. Rowe. “Sperling 559 RA, Cummings JL (2015) Alzheimer’s disease.” Nat. Rev. Dis. Primers 1 (2015): 15056.

[2] Gauthier, Serge, Barry Reisberg, Michael Zaudig, Ronald C. Petersen, Karen Ritchie, Karl Broich, Sylvie Belleville et al. “Mild cognitive impairment.” The lancet 367, no. 9518 (2006): 1262–1270.

[3] Winblad, Berndt, Katie Palmer, Miia Kivipelto, Vesna Jelic, Laura Fratiglioni, L-O. Wahlund, Agneta Nordberg et al. “Mild cognitive impairment–beyond controversies, towards a consensus: report of the International Working Group on Mild Cognitive Impairment.” Journal of internal medicine 256, no. 3 (2004): 240–246.

[4] Bruscoli, Maddalena, and Simon Lovestone. “Is MCI really just early dementia? A systematic review of conversion studies.” International Psychogeriatrics 16, no. 2 (2004): 129.

[5] Marcus, Daniel S., Tracy H. Wang, Jamie Parker, John G. Csernansky, John C. Morris, and Randy L. Buckner. “Open Access Series of Imaging Studies (OASIS): cross-sectional MRI data in young, middle aged, nondemented, and demented older adults.” Journal of cognitive neuroscience 19, no. 9 (2007): 1498–1507.

[6] Pangman, Verna C., Jeff Sloan, and Lorna Guse. “An examination of psychometric properties of the mini-mental state examination and the standardized mini-mental state examination: implications for clinical practice.” Applied Nursing Research 13, no. 4 (2000): 209–213.

[7] Zanaty, E. A. “Support vector machines (SVMs) versus multilayer perception (MLP) in data classification.” Egyptian Informatics Journal 13, no. 3 (2012): 177–183.

[8] Gardner, Matt W., and S. R. Dorling. “Artificial neural networks (the multilayer perceptron)—a review of applications in the atmospheric sciences.” Atmospheric environment 32, no. 14-15 (1998): 2627–2636.

[9] Hornik, Kurt, Maxwell Stinchcombe, and Halbert White. “Multilayer feedforward networks are universal approximators.” Neural networks 2, no. 5 (1989): 359–366.

[10] Chawla, N. V., Bowyer, K. W., Hall, L. O., & Kegelmeyer, W. P. (2002). SMOTE: synthetic minority over-sampling technique. Journal of artificial intelligence research, 16, 321–357.

[11] Saito, Takaya, and Marc Rehmsmeier. “The precision-recall plot is more informative than the ROC plot when evaluating binary classifiers on imbalanced datasets.” PloS one 10, no. 3 (2015): e0118432.

